# Efficient Dlx2-mediated astrocyte-to-neuron conversion and inhibition of neuroinflammation by NeuroD1

**DOI:** 10.1101/2022.07.11.499522

**Authors:** Min-Hui Liu, Yu-Ge Xu, Xiao-Ni Bai, Jian-Hua Lin, Zong-Qin Xiang, Tao Wang, Liang Xu, Wen Li, Gong Chen

## Abstract

*In vivo* astrocyte-to-neuron (AtN) conversion induced by overexpression of neural transcriptional factors has great potential for neural regeneration and repair. Here, we demonstrate that a single neural transcriptional factor Dlx2 converts mouse striatal astrocytes into neurons in a dose-dependent manner. Lineage-tracing studies in Aldh1l1-CreER^T2^ mice confirm that Dlx2 can convert striatal astrocytes into DARPP32^+^ and Ctip2^+^ medium spiny neurons (MSNs). Time-course studies reveal a gradual conversion from astrocytes to neurons in 1 month, with a distinct intermediate state in-between astrocytes and neurons. Interestingly, when Dlx2-infected astrocytes start to lose astrocytic markers, the other local astrocytes proliferate to maintain astrocytic level in the converted areas. Unexpectedly, while Dlx2 efficiently reprograms astrocytes into neurons in the grey matter striatum, it also induces partial reprogramming of astrocytes in the white matter corpus callosum. Such partial reprogramming of white matter astrocytes is associated with neuroinflammation, which can be essentially suppressed by the addition of NeuroD1. Our results highlight the importance of investigating AtN conversion both in the grey matter and white matter in order to thoroughly evaluate therapeutic potentials. This study also unveils a critical role of anti-inflammation by NeuroD1 during AtN conversion.

## Introduction

Endogenous adult neurogenesis is limited in the mammalian central nervous system (CNS), making it difficult to self-repair after neural injury or diseases. The advent of induced pluripotent stem cell technology ^1,2^ provides a potential cell-replacing therapy, but also faces potential risks of immune rejection and tumorigenesis ^3^. Alternatively, manipulation of endogenous cells to generate new neurons in the adult brain and spinal cord has been investigated as a new approach for neural repair ^4–6^. Glial cells in the adult brain can be activated after injury to proliferate and dedifferentiate into neuroblasts through the inhibition of Notch signaling ^7,8^. A more efficient way of neuroregeneration is to convert reactive glial cells into functional new neurons via overexpression of neurogenic transcription factors (TFs; NeuroD1, Ascl1, Sox2, Neurogenin2, etc.) ^9–17^ or down-regulation of RNA-binding protein PTBP1 ^18,19^. This *in vivo* glia-to-neuron conversion strategy has been reported to promote functional recovery in a series of rodent models including ischemic stroke, Huntington’s disease, Parkinson’s disease, spinal cord injury and epilepsy ^18–25^. Our recent work in the non-human primate brain further demonstrate that *in vivo* glia-to-neuron conversion can repair neural tissue after ischemic insult ^26^. These results suggest that *in vivo* glia-to-neuron conversion may be developed into a potential new therapy for neural repair.

On the other hand, a few studies raised concerns on *in vivo* glia-to-neuron conversion based on a special lineage-tracing transgenic mice ^27–29^. We have previously reported that microglia cannot be converted into neurons because microglia originates from hematopoietic stem cells not neural stem cells (Guo et al., 2014; Supplementary data) ^9^. PTBP1 is quite controversial at this moment and requires more studies to clarify ^18,19,28,30^. On NeuroD1-mediated astrocyte-to-neuron conversion, following our initial study and a series of continuous studies ^9,20,23,24,26,31^, many other labs have successfully repeated our NeuroD1 conversion results ^32–35^. Moreover, our lab and Davidson lab have used the same astrocytic lineage-tracing mice (Aldh1|1-CreER^T2^) to demonstrate that NeuroD1 can convert lineage-traced astrocytes into neurons, although conversion efficiency varies in different studies ^14,36,37^. Importantly, we show that lineage-traced astrocytes are more difficult to convert and require enhancer to induce higher expression level of NeuroD1 to overcome the conversion barrier ^37^.

Regeneration of proper neuronal subtypes in specific brain regions is one of the critical requirements for functional recovery. Previous studies have found that the subtypes of converted cells are determined by both transcription factors and local environment. NeuroD1 and Neurogenin2 play important role in the development of cortical excitatory neurons, and are reported to convert glial cells into glutamatergic neurons ^9,13^. Ascl1 has been reported to convert midbrain astrocytes into both glutamatergic and GABAergic neurons ^10^. NeuroD1+Ascl1+Lmx1a+miR218 has been reported to reprogram striatal astrocytes into dopaminergic neurons ^38^. We have previously reported that NeuroD1+Dlx2 converts striatal astrocytes into GABAergic neurons and significantly extended the life span of R6/2 mouse model of Huntington’s disease ^21^. Since Dlx2 is a homeobox transcription factor that is critical for the development of striatal neurons ^39^, we carry out further investigation on Dlx2 alone following our earlier studies.

In this study, we ectopically expressed Dlx2 in mouse striatal astrocytes and used different virus systems to investigate the effect of Dlx2 alone on astrocytes. We first transduced retrovirus-Dlx2 to the stab-injured mouse striatum and detected both DCX^+^ immature neurons and NeuN^+^ neurons among Dlx2-infected cells. Next, we used astrocytic lineage-tracing mice (Aldh1|1::CreER^T2^) to express Dlx2 specifically in astrocytes and found efficient generation of striatal neurons (DARPP32^+^ and Ctip2^+^) originated from lineage-traced astrocytes. We further employed GFAP::Dlx2-GFP to conduct extensive dose-finding and time course studies and discovered a clear intermediate state in-between astrocytes and neurons during AtN conversion process. A surprising finding was the detection of unexpected neural inflammation in the corpus callosum following Dlx2 infection, which was alleviated after the addition of NeuroD1. These results suggest that Dlx2 alone is sufficient to induce AtN conversion in the striatum but may result in neural inflammation in the white matter. Our studies also reveal an important role of NeuroD1 on antiinflammation during neuronal conversion.

## Result

### Retroviral expression of Dlx2 converts reactive glial cells into neurons in injured striatum

Our previous studies have demonstrated that NeuroD1 plus Dlx2 can convert striatal astrocytes into medium spiny neurons (MSN) both in wildtype mice and R6/2 or YAC128 transgenic mouse models of Huntington’s disease ^21^. Since NeuroD1 is not involved in striatal development ^40,41^, in this study we removed NeuroD1 and investigated the effect of Dlx2 alone on striatal glial cell conversion. To unambiguously target dividing reactive glial cells rather than non-dividing postmitotic neurons, we first injected retroviruses (CAG::GFP-IRES-GFP or CAG::Dlx2-IRES-GFP) that specifically expressed ectopic genes in dividing glial cells induced by stab-injury in the mouse striatum (Fig. 1A for design). As expected, many of the control retrovirus GFP-infected cells were Sox9^+^ glial cells at 7 days post infection (7 D, Fig. 1B). At 30 days post-viral injection (30 D), the retroviral infected cells (GFP^+^) in the control group were a mixture of glial cells, including astrocytes (Sox9^+^), oligodendrocytes or its precursor cells (Olig2^+^), and microglia (Iba1^+^) (Fig. 1B and Fig. S1C, quantified in Fig. 1D). Neuroblasts/immature neurons (DCX^+^) or mature neurons (NeuN^+^) were rarely detected in the control group infected by GFP retrovirus (Fig. 1C-D, Fig. S1A and S1C). These results indicate that retroviruses only infect dividing glial cells but not non-dividing neurons in the stab-injured striatum.

**Figure 1.**
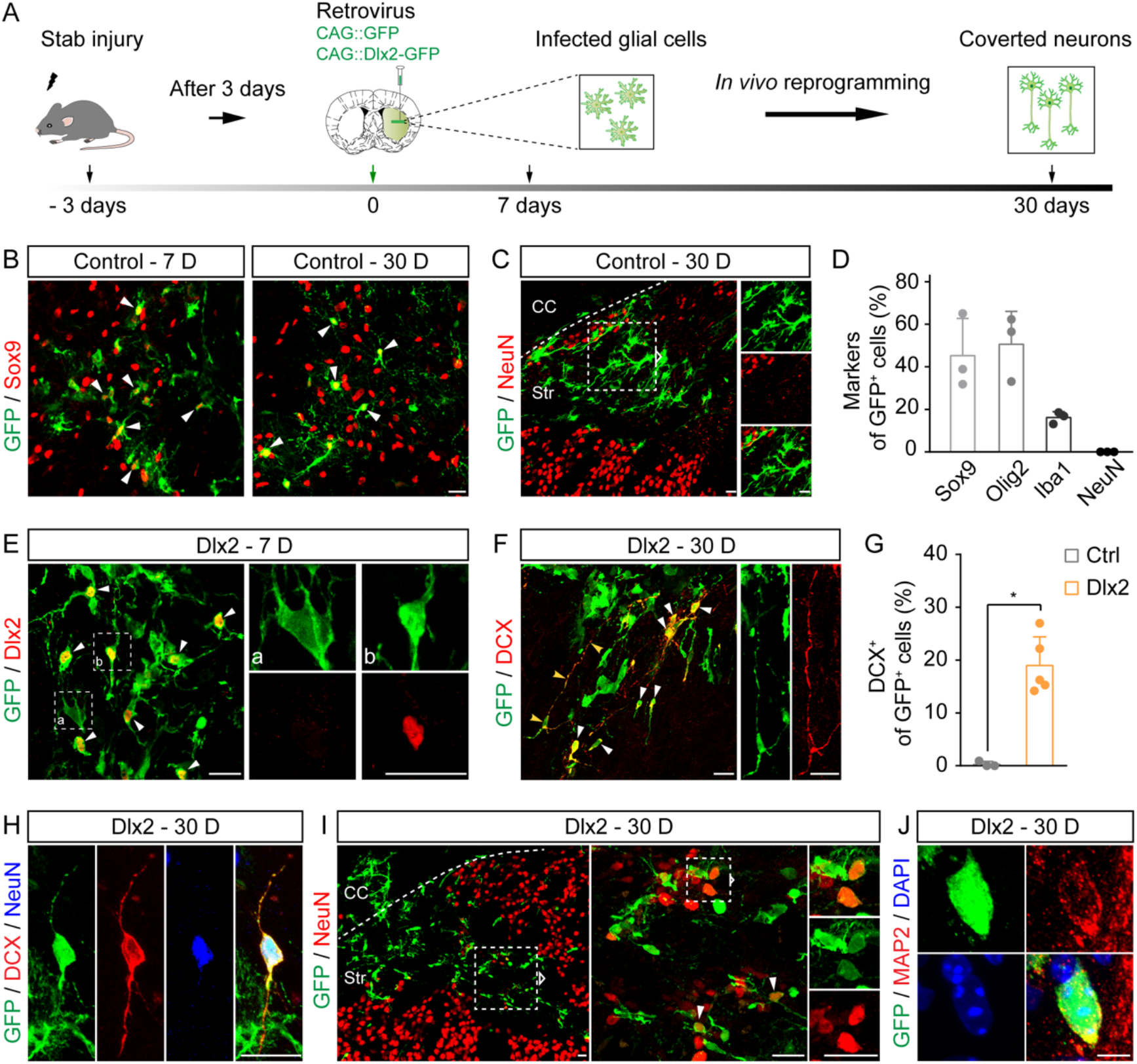
Retroviral expression of Dlx2 reprograms striatal glia into neurons after stab injury. (A) Experimental design to investigate striatal glia-to-neuron conversion through the retrovirus system after stab injury. (B) Control retrovirus GFP-infected cells (green) consistently express astrocytic marker Sox9 (red) from 7 to 30 days post viral infection. Arrowheads indicate some GFP and Sox9 co-labeled cells. (C) None of the control retrovirus GFP-infected cells express NeuN (red). Dash line to distinguish corpus callosum (CC) and striatum (Str). (D) Quantification of control virus GFP-infected cells expressing glial markers (Sox9, 45.3 ± 17.5%; Olig2, 50.6 ± 15.5%; Iba1, 16.2 ± 2.8%; mean ± SD; n = 3 animals) or neuronal marker (NeuN, not detected) at 30 days post viral injection. (E) Representative images confirming the ectopic expression of Dlx2 (red) in the retrovirus Dlx2-GFP-infected cells at 7 D (white arrowheads indicate the soma, and dotted boxes indicate different cell morphology that is enlarged in the right two images). (F) At 30 days after the retrovirus-Dlx2 infection, immature neuronal marker DCX (red) was detected among Dlx2-GFP-infected cells (white arrowheads indicate the soma, and yellow arrowheads indicate the neurite-like process that is enlarged in the right two images). (G) Quantified data showing a considerable portion of DCX positive cells generated in the Dlx2 group, while in the control group the percentage is quite low (Ctrl, 0.3 ± 0.5%, n = 3 animals; Dlx2, 19.0 ± 5.4%, n = 5 animals; mean ± SD; *\*P* < 0.05, Mann Whitney test). (H) Representative images show that some DCX (red) and NeuN (blue) double positive cells are also observed in Dlx2-GFP-infected cells. (I and J) Dlx2-infected cells (green) express neuronal markers NeuN (I, red) or MAP2 (J, red). Scale bars: 20 μm, except 5 μm for J.

After confirming the infection of dividing glial cells, we injected retrovirus Dlx2-GFP into the stab-injured striatum. At 7 days after virus infection (7 D), the expression of ectopic Dlx2 was clearly detected in the infected cells (Fig. 1E). Interestingly, at this early stage at 7 D, while some Dlx2-GFP-infected cells remained Sox9^+^ astrocytes (Fig. S1B, rarely NeuN^+^ signal), some other Dlx2-infected cells had changed from astrocytic morphology into bipolar morphology with thin neurite-like processes (Fig. 1E, see enlarged panel b). By 30 days after Dlx2 infection, ~20% of Dlx2-infected cells became DCX^+^ immature neurons, but in the control group DCX^+^ cells were rarely detected (Fig. 1F-G, Fig. S1C). Among these Dlx2-induced DCX^+^GFP^+^ cells, some of them extended long neurite-like structure (Fig. 1F and 1H) and even co-expressed neuronal marker NeuN (Fig. 1H), suggesting a transition from immature to mature neurons after 30 days of Dlx2 expression. Furthermore, there was a small portion of Dlx2-GFP-infected cells immunopositive for mature neuronal markers of both NeuN and MAP2 (Fig. 1I and 1J). Together, these data indicate that Dlx2 alone can reprogram proliferative glial cells into neurons in the injured striatum *in vivo*.

### Lineage-tracing studies reveal Dlx2-converted neurons originated from astrocytes

Since the retrovirus system targeted not only dividing astrocytes but also other glial cells, we further investigated whether Dlx2 alone can convert astrocytes *per se* into neurons. To answer this question, we used astrocytic lineage-tracing mice Aldh1l1-CreER^T2 42^ to trace the origin of converted neurons. Tamoxifen was administered to induce Cre-mediated recombination after injecting control AAV9 Flex-mCherry-P2A-mCherry or AAV9 Flex-Dlx2-P2A-mCherry into the striatum of Aldh1l1-CreER^T2^ mice (Fig. 2A). As expected, the control virus mCherry specifically labeled astrocytes in the striatum of Aldh1l1::CreER^T2^mice, as shown by ~93% of mCherry^+^ cells coexpressing astrocytic markers S100β and GFAP (Fig. 2B-C). In the meantime, we did not detect Olig2^+^ or Iba1^+^ nor NeuN^+^ cells among control AAV9 Flex-mCherry-infected cells (Fig. S2A-B, Fig. 2C and 2E-F), suggesting that this Aldh1l1::CreER^T2^ + AAV9 Flex-mCherry system is specific in tracing the lineage of astrocytes.

**Figure 2.**
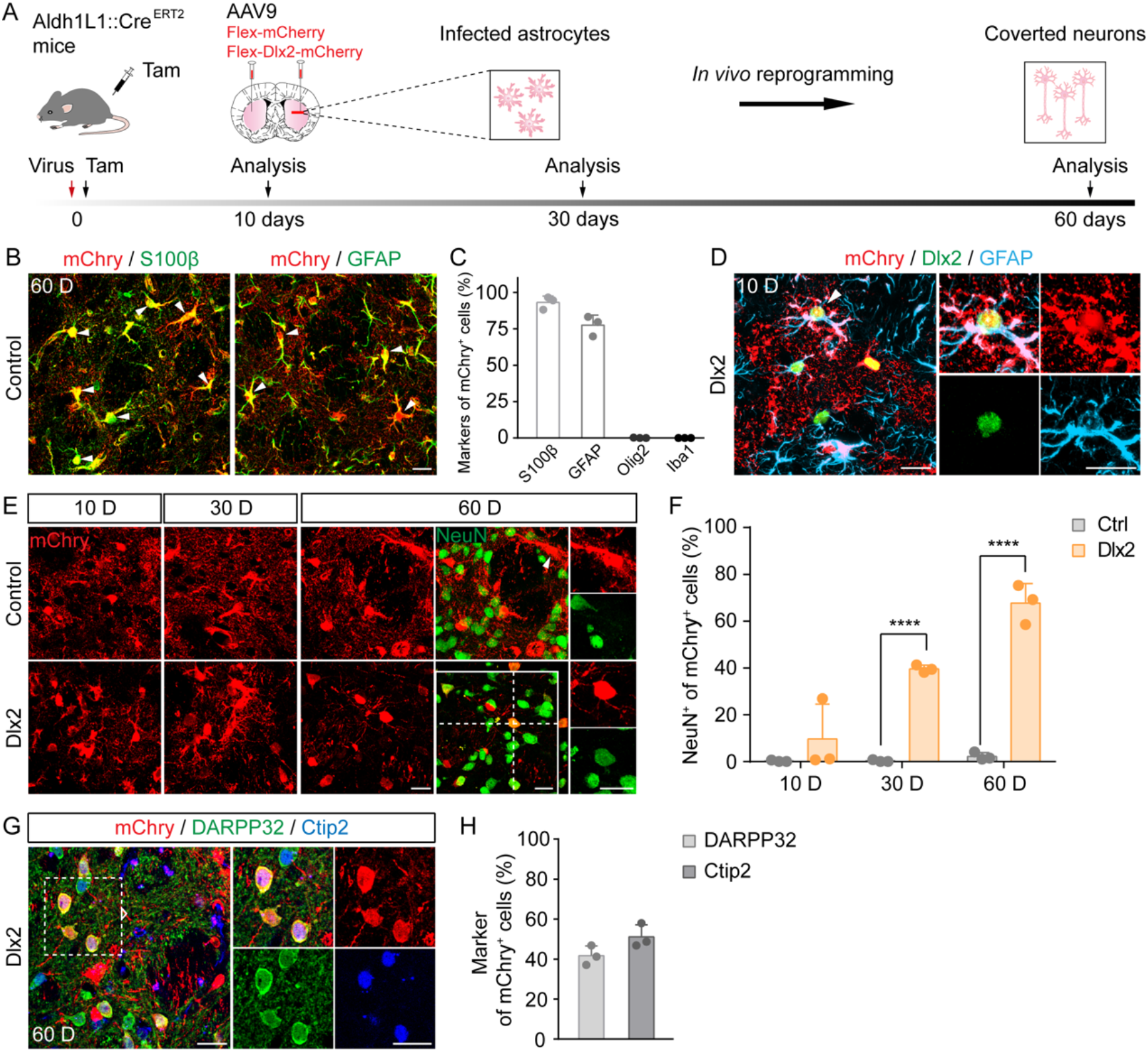
Striatal astrocytes are converted into medium spiny neurons in lineage-tracing Aldh1l1::CreER^T2^ mice. (A) Experimental design to investigate striatal astrocyte-to-neuron conversion in lineage-tracing Aldh1l1::CreER^T2^ mice using AAV9 FLEX system. (B) Representative images show that the control virus (AAV9 Flex-mCherry; red) are specifically expressed in astrocytes (S100β; GFAP; green) at 60 days. Arrowheads indicate astrocytic markers and mCherry co-labeled cells. (C) Quantitation showing the majority of the control virus-infected cells are positive for astrocytic markers (S100β, 93.1 ± 4.4%; GFAP, 77.6 ± 6.9%; Olig2, 0.1 ± 0.25%; Iba1, not detected; mean ± SD; n = 3 animals), confirming the specificity of astrocyte lineage-tracing in Aldh1l1::CreER^T2^ mice. (D) Representative images show the co-expression of Dlx2 (green) and GFAP (blue) in the mCherry^+^ cells (red) after 10 days of injecting Dlx2-mCherry into the mouse striatum. The arrowhead indicates the co-localization of mCherry^+^cells together with Dlx2 and GFAP. (E) Representative images show that Dlx2-infected cells (bottom row, mCherry^+^, red) change from astrocyte-like morphology at 10 D to neuron-like morphology at 60 D, while those in the control group (top panel) remain astrocyte-like morphology. Enlarged images display no NeuN signal (arrowhead) detected in the the control mCherry-infected cells, while most Dlx2-mCherry-infected cells are co-labeled with NeuN signal (green) at 60 days after viral injection. (F) Quantified data showing a gradual increase in the percentage of NeuN^+^cells among Dlx2-mCherry-infected cells at different conversion time points, while the NeuN^+^ rate remains at quite low in the control mCherry group (Ctrl group: 10 D, 0.2 ± 0.4%; 30 D, 0.3 ± 0.5%; 60 D, 2.2 ± 1.7%. Dlx2 group: 10 D, 9.6 ± 15.0%; 30 D, 39.6 ± 1.6%; 60 D, 67.7 ± 8.4%. mean ± SD. *\*\*\*\*P* < 0.0001, two-way ANOVA with Sidak’s multiple comparison test. n = 3 animals). (G) Representative images show that the Dlx2-converted neurons (red) express the markers for the medium spiny neurons (MSN; DARPP32, green; Ctip2, blue). (H) Quantification of converted MSNs (DARPP32, 41.7 ± 4.9%; Ctip2, 51.2 ± 6.0%; mean ± SD; n = 3 animals). Scale bars: 20 μm.

Next, we examined the effect of Dlx2 on the lineage-traced astrocytes. Similar to our retroviral expression of Dlx2, we first confirmed Dlx2 expression in the lineage-traced mCherry^+^ astrocytes at 10 days after virus infection (Fig. 2D). Side-by-side comparison between the control and Dlx2 group at different time points revealed clear difference in morphological changes among the lineage-traced astrocytes. In the control group, mCherry-traced astrocytes did not change their morphology from 10 D to 60 D (Fig. 2E, top row). In contrast, most Dlx2-mCherry-infected cells were initially in astrocytic morphology with many branches at 10 D but later transformed into neuron-like morphology with a few processes by 60 days after Dlx2 expression (Fig. 2E, bottom row). Quantitatively, NeuN immunostaining revealed a time-dependent conversion of mCherry^+^ astrocytes into mCherry^+^ neurons in the Dlx2 group (Fig. 2F). While the control mCherry-infected cells rarely showed NeuN^+^ signal, the number of NeuN^+^ cells among Dlx2-mCherry-infected ones reached ~70% at 60 D (Fig. 2E-F). We further analyzed the subtypes of the newly generated neurons originated from mCherry^+^ astrocytes, and found that ~40-50% of the Dlx2-converted neurons were DARPP32^+^ and Ctip2^+^ medium spiny neurons (MSNs) (Fig. 2G-H). These results suggest that lineage-traced striatal astrocytes can be reprogrammed into striatal MSNs after ectopic expression of Dlx2.

### Dose-dependent conversion by Dlx2

We next investigated what is an appropriate dose for Dlx2 to trigger AtN conversion by using astrocyte-specific GFAP promoter to drive ectopic gene expression (GFAP::GFP as the control group and GFAP::Dlx2-P2A-GFP as the experimental group, Fig. 3A). Previous study has reported neurotoxicity induced by high titer of AAV ^43^. Therefore, we tested a series of different dosages (2E10, 2E11, 2E12, 1E13 GC/ml, virus volume fixed at 2 μl for all titres) of control AAV9 GFAP::GFP in the mouse striatum. After conducting Iba1 immunostaining to assess microglia activation in the striatum, we observed a severe inflammatory response when the virus titer reached 1E13 GC/mL, whereas microglia appeared to be normal at lower titre of 2E10 to 2E12 GC/mL (Fig. 3B). This control experiment informed us that AAV titre should not reach 1E13 GC/ml when injected into the parenchyma of the mammalian brains. Otherwise, severe neural inflammation may cause unwanted side effects and make data interpretation complicated or impossible.

**Figure 3.**
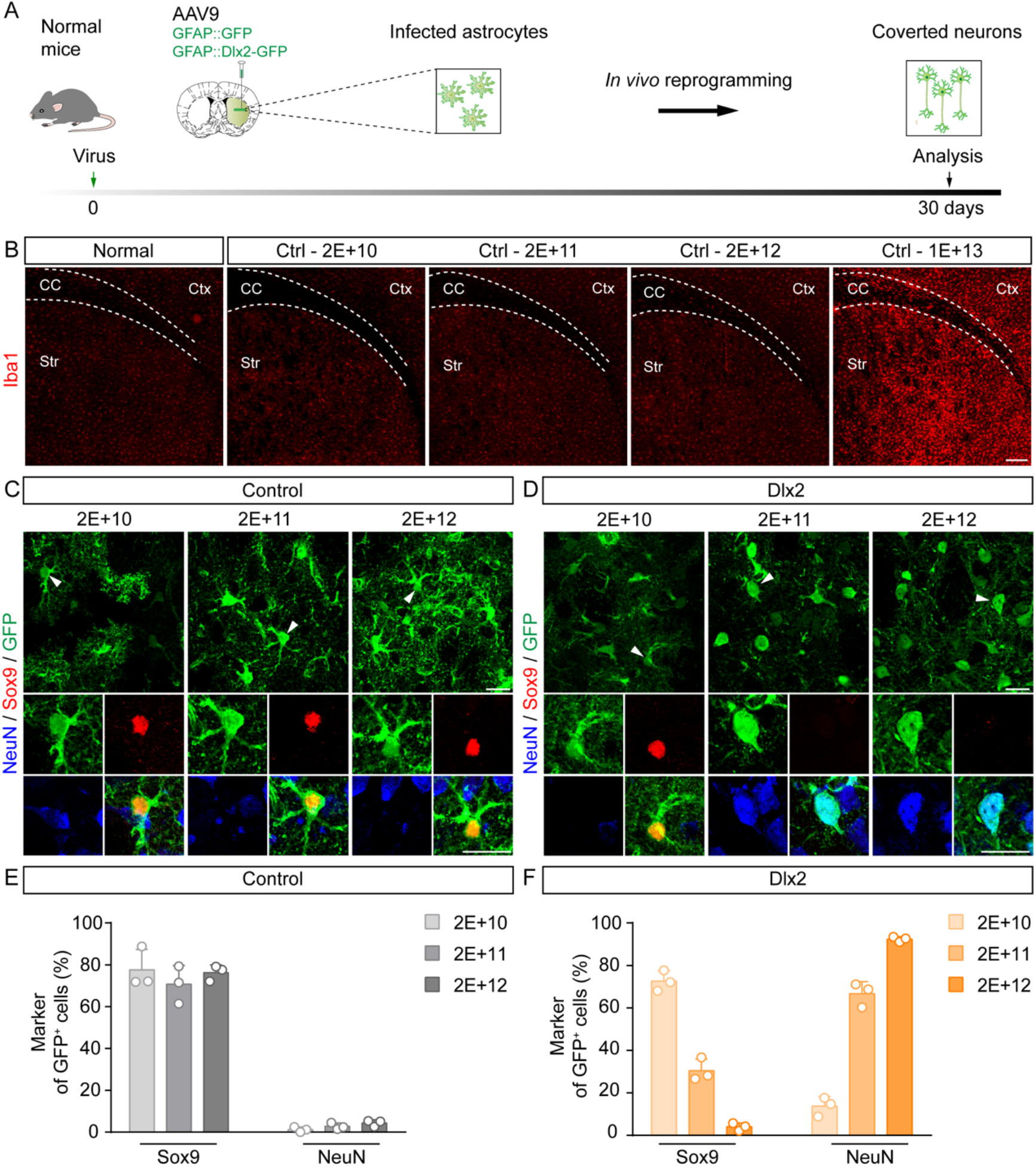
Dlx2-induced astrocyte-to-neuron conversion exhibits a dose-dependent manner. (A) Experimental design to investigate striatal AtN conversion using AAV9 GFAP promoter system. (B) Representative images reveal that high titer (1E13 GC/mL, 2 μl) of AAV9 GFAP::GFP causes Iba1^+^ microglia (red) activation in the striatum (Str, striatum; Ctx, cortex; CC, corpus callosum). (C) Representative images show that the infected cells (green) in the control group at all virus titers tested specifically express astrocytic marker Sox9 (red, arrowhead). NeuN^+^ signal (blue) is rarely detected among the control virus GFP-infected cells. (D) Representative images showing the morphological changes of Dlx2-infected cells (green) among different viral dosages. Note that NeuN (blue) signal can be detected among higher titre of Dlx2-infected cells at 2E11 and 2E12, but not at low titre of 2E10. (E) Quantified data shows that the control virus GFP-infected cells mainly express astrocytic marker Sox9 at all three different titres (Titre: 2E+10 GC/mL, 77.6 ± 9.7%; 2E+11 GC/mL, 70.7 ± 9.0%; 2E+12 GC/mL, 76.3 ± 3.7%; mean ± SD; n = 3 animals), but rarely express neuronal marker NeuN (Titre: 2E+10 GC/mL, 1.2 ± 1.2%; 2E+11 GC/mL, 2.8 ± 1.6%; 2E+12 GC/mL, 4.4 ± 1.6%; mean ± SD; n = 3 animals). (F) Quantification shows that conversion rate of Dlx2 increases in a dose-dependent manner (Sox9: 2E+10 GC/mL, 72.6 ± 4.9%; 2E+11 GC/mL, 30.5 ± 5.5%; 2E+12 GC/mL, 4.0 ± 1.8%. NeuN: 2E+10 GC/mL, 13.8 ± 4.5%; 2E+11 GC/mL, 66.7 ± 5.7%; 2E+12 GC/mL, 92.4 ± 1.2%. mean ± SD. *\*P* < 0.05, *\*\*P* < 0.01, *\*\*\*P* < 0.001, two-way ANOVA with Sidak’s multiple comparison test. n = 3 animals). Scale bars: 200 μm for B, 20 μm for C-D.

Based on the dose-finding studies using control AAV9 GFAP::GFP, we next investigated the effect of Dlx2 on AtN conversion using a relatively safe range of dosing (2E10, 2E11, and 2E12 GC/ml; all in 2 μl). The same dosing of control AAV9 GFAP::GFP was used for side-by-side comparison. After 30 days of viral injection, the control GFAP::GFP-infected cells consistently displayed typical astrocytic morphology and expressed astrocytic marker Sox9 at all three different titres (Fig. 3C; 2E10, 2E11, 2E12 GC/ml; 2 μl). In contrast, for Dlx2 group, while most cells infected by low dose (2E10 GC/ml) of Dlx2 displayed astrocytic morphology, higher dose of Dlx2 (2E11 and 2E12 GC/ml) induced morphological change into neurons with NeuN expression at 30 D (Fig. 3D). Quantitatively, ~70-80% of control GFAP::GFP-infected cells were Sox9^+^astrocytes at different titres from 2E10 to 2E12 GC/ml, and only <5% of GFAP::GFP-infected cells showed NeuN^+^ signal at high dose of 2E12 GC/ml (Fig. 3E, 30 D). Therefore, this range of doses (2E10 to 2E12 GC/ml, 2 μl) has minimal leakage of transgene expression in neurons due to the restriction of astrocytic promoter GFAP. In contrast to the control group, quantitation of the Dlx2 group revealed a clear dose-dependent changes among the Dlx2-infected cells (Fig. 3F). The majority of cells infected by low dose Dlx2 (2E10 GC/ml) remained Sox9^+^ astrocytes at 30 D, whereas high dose (2E12 GC/ml) Dlx2-infected cells had converted into NeuN^+^ neurons (Fig. 3F; conversion efficiency: 2E10 GC/ml, 14%; 2E11 GC/ml, 67%; 2E12 GC/ml, 92%). Together, these experiments demonstrate that Dlx2-mediated AtN conversion is dose-dependent.

### Time-dependence of Dlx2-induced AtN conversion

After determining the dose-dependence of cell conversion, we further investigated the time-dependence of AtN conversion induced by Dlx2 at a fixed dose (2E12 GC/ml, 2 μl). We performed analyses at a series of different time points, including 2, 5, 10, 15, 20, and 30 days after Dlx2 infection, and made several interesting findings. At early stage such as 2 days after Dlx2 infection, the majority of AAV9-Dlx2 infected cells displayed astrocytic morphology and expressed astrocyte marker Sox9 (Fig. 4B, left column). However, at later stage such as 30 days after Dlx2 infection, the majority of Dlx2-infected cells became NeuN^+^ neurons (Fig. 4B, right column). Interestingly, during first two weeks of conversion, some Dlx2-infected cells had lost astrocytic marker Sox9 but not yet acquired neuronal marker NeuN (Fig. 4B, middle column), suggesting that they are neither astrocytes nor neurons. We termed this stage as an intermediate state during AtN conversion, which rarely occurs in the normal brain. Importantly, we examined whether these intermediate state cells would undergo apoptosis by performing TUNEL staining. The results showed that except a few TUNEL^+^ signal in the needle track due to stab injury caused by viral injection, there were no apoptotic cells detected among the Dlx2-GFP-infected cells (Fig. S3), excluding the cell death hypothesis during conversion. Such intermediate state cells gradually decreased after the completion of AtN, such as after 30 days of Dlx2 infection (Fig. 4C, green bars). Quantitative analyses revealed a clear decrease of Sox9^+^ astrocytes among Dlx2-infected cells from 80% at 2 D to 4% at 30 D (Fig. 4C, blue bars), with a concomitant increase of NeuN^+^ cells from 7% at 2 D to 92% at 30 D (Fig. 4C, brown bars). Such simultaneous changes from astrocytic population to neuronal population among Dlx2-infected cells suggest a clear AtN conversion in the mouse striatum. The green bars in Fig. 4C (15-20%) indicate a transitional stage (both Sox9^-^ and NeuN^-^) in-between astrocytes and neurons that were mainly detected at early stage but rarely at later stage of AtN conversion. These results demonstrate that Dlx2 converts striatal astrocytes into neurons in a time-dependent manner, with a distinct intermediate state in-between astrocytes and neurons during the conversion process.

**Figure 4.**
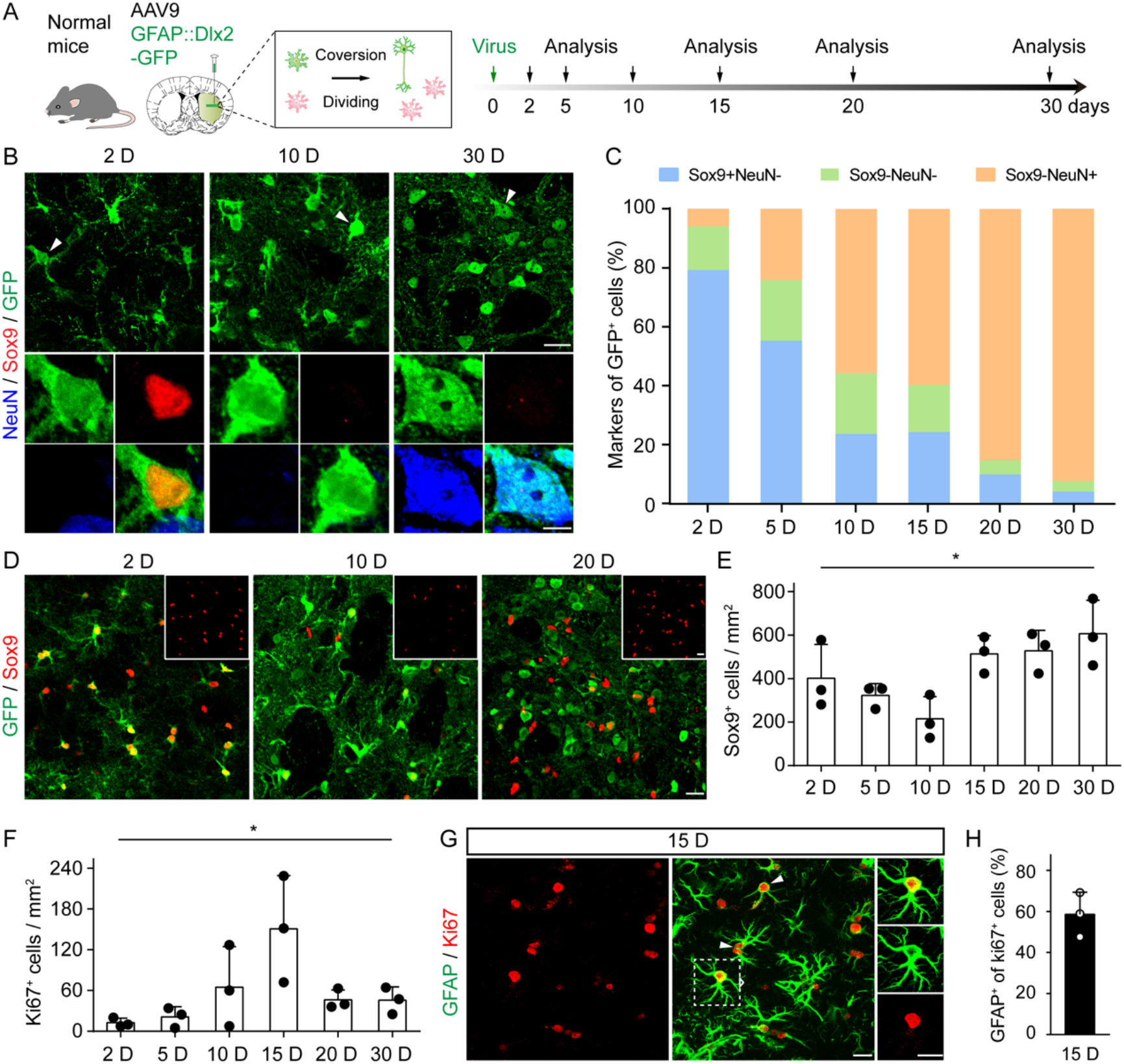
Striatal astrocytes replenish themselves following astrocyte-to-neuron conversion. (A) Experimental design to trace striatal astrocyte-to-neuron conversion at different time points. (B) Representative images showing the gradual morphological change of Dlx2-infected cells (green, arrowheads) as the conversion proceeds, which is accompanied with a gradual loss of astrocytic marker (Sox9, red) and gain of neuronal marker (NeuN, blue). There is an intermediate state featured by Sox9^-^NeuN^-^ of the Dlx2-infected cells around 10 D. (C) Quantification of the three types of cells (Sox9^+^NeuN^-^; Sox9^-^NeuN^-^; Sox9^-^NeuN^+^) among the Dlx2-infected cells at different time points of AtN conversion. (Sox9^+^NeuN^-^: 2 D, 79.2%; 5 D, 55.3%; 10 D, 23.6%; 15 D, 24.3%; 20 D, 9.9%; 30 D, 4.0%. Sox9^-^NeuN^-^: 2 D, 14.8%; 5 D, 20.6%; 10 D, 20.7%; 15 D, 16.0%; 20 D, 4.8%; 30 D, 3.6%. Sox9^-^NeuN^+^: 2 D, 7.1%; 5 D, 24.2%; 10 D, 55.7%; 15 D, 59.7%; 20 D, 85.3%; 30 D, 92.4%. n = 3 animals for each time point). (D and E) Representative images (D) and quantification (E) showing an initial decrease of Sox9 signal (red) followed by a quick rebound within the Dlx2-infected areas (GFP, green). The inlets show the Sox9 signal only. Quantified data in Panel E: 2 D, 402 ± 156; 5 D, 323 ± 54; 10 D, 216 ± 102; 15 D, 513 ± 85; 20 D, 528 ± 94; 30 D, 607 ± 154. Mean ± SD. *\*P* < 0.05, one-way ANOVA test. n = 3 animals for each time point). (F) Quantification showing the change of Ki67^+^ proliferative cells at different time points within the infected areas of mouse striatum in the Dlx2 group. Note an increase in the number of proliferative cells (Ki67) at 15 D: 2 D, 13 ±7; 5 D, 21 ± 15; 10 D, 65 ± 60; 15 D, 151 ± 78; 20 D, 46 ± 15; 30 D 46 ± 20. *\*P* < 0.05, one-way ANOVA test. n = 3 animals). (G and H) Representative images (G) and quantification (H) show that most Ki67^+^ cells (red) are co-labeled with astrocytic marker GFAP (green). Enlarged images indicate one co-labeled cell. (H, 58.6 ± 10.8, mean ± SD, n = 3 animals). Scale bars: 20 μm, except B bottom panels for 5 μm.

### Astrocytes replenish themselves following AtN conversion

Astrocytes are one of the most prevalent types of glia and play indispensable roles in maintaining the homeostasis in the CNS ^44^. If astrocytes are converted into neurons, is there a potential side effect associated with astrocyte conversion? To answer this question, we analyzed the density of astrocytes at different time points during AtN conversion (Fig. 4A for experimental design). After injecting the AAV9 GFAP::Dlx2-GFP viruses, we observed a transient decrease in the number of Sox9^+^ astrocytes around 10 days, suggesting that some astrocytes were converted into neurons and hence lost astrocytic marker Sox9 (Fig. 4D-E). The Dlx2-induced decrease of Sox9^+^ cells was coupled with an increase of NeuN^+^ cells around 10 days (Fig. 4C, brown bar at 10 D), further confirming a cell fate change from astrocytes to neurons. Such Dlx2-induced AtN conversion showed no adversary effect on local striatal neurons (Fig. S4A). Interestingly, the number of Sox9^+^ astrocytes only decreased transiently around 10 D, followed by a quick rebound by 15 days after Dlx2 infection (Fig. 4E), suggesting that remaining astrocytes might have proliferated to maintain their homeostasis following conversion. To investigate this possibility, we employed Ki67, a marker labeling proliferative cells, to examine cell proliferation during AtN conversion. Strikingly, we found a significant increase in the number of Ki67^+^ cells from 10 to 15 days after Dlx2 infection (Fig. 4F and S4B), which coincided well with the transient decrease and quick rebound of the Sox9^+^astrocytes (Fig. 4E). Indeed, coimmunostaining of Ki67 and astrocytic marker GFAP at 15 D confirmed that the majority of Ki67^+^ cells (~60%) were GFAP^+^astrocytes (Fig. 4G-H). These results indicate that accompanying the AtN conversion, local astrocytes can proliferate and replenish themselves.

### Dlx2 induces inflammation in the white matter

During our investigation of Dlx2-induced AtN conversion, we noticed a significant activation of microglia in the axon bundles of the striatum as well as the white matter of corpus callosum (Fig. 5). Such activated microglia were not detected in the control group (Fig. S5A-C). Since activation of microglia might be associated with neuroinflammation, we were alerted by this unexpected phenomenon and conducted further investigation. We analyzed a series of time points (Fig. 5A) and observed a time-dependent activation of microglia in the striatum infected by Dlx2, which increased at 10 D, persisted to 30 D, and then declined at 60 D (Fig. 5B-C, Str, striatum). Besides the striatum, we also found an increase of Iba1 signal in the corpus callosum infected by Dlx2, which started around 10 D and became even more intense at 60 D (Fig. 5B-C, CC, corpus callosum). In addition to Iba1, we further examined other neuroinflammation markers including CD68 (marker for phagocytic microglia and macrophages) ^45^ and CD45 (marker for infiltrating monocytes) ^46^. Consistent with the increase of Iba1 signal, compared to the control virus (Fig. 5E left column, and Fig. S6 for high power images), both CD68 and CD45 signal increased significantly in the corpus callosum infected by Dlx2 at 60 D (Fig. 5E middle column, and Fig. S6). This unexpected finding of Dlx2-induced neuroinflammation suggests that AtN conversion shall be carefully analyzed at different doses and different time points to evaluate its potential side effects in both grey matter and white matter.

**Figure 5.**
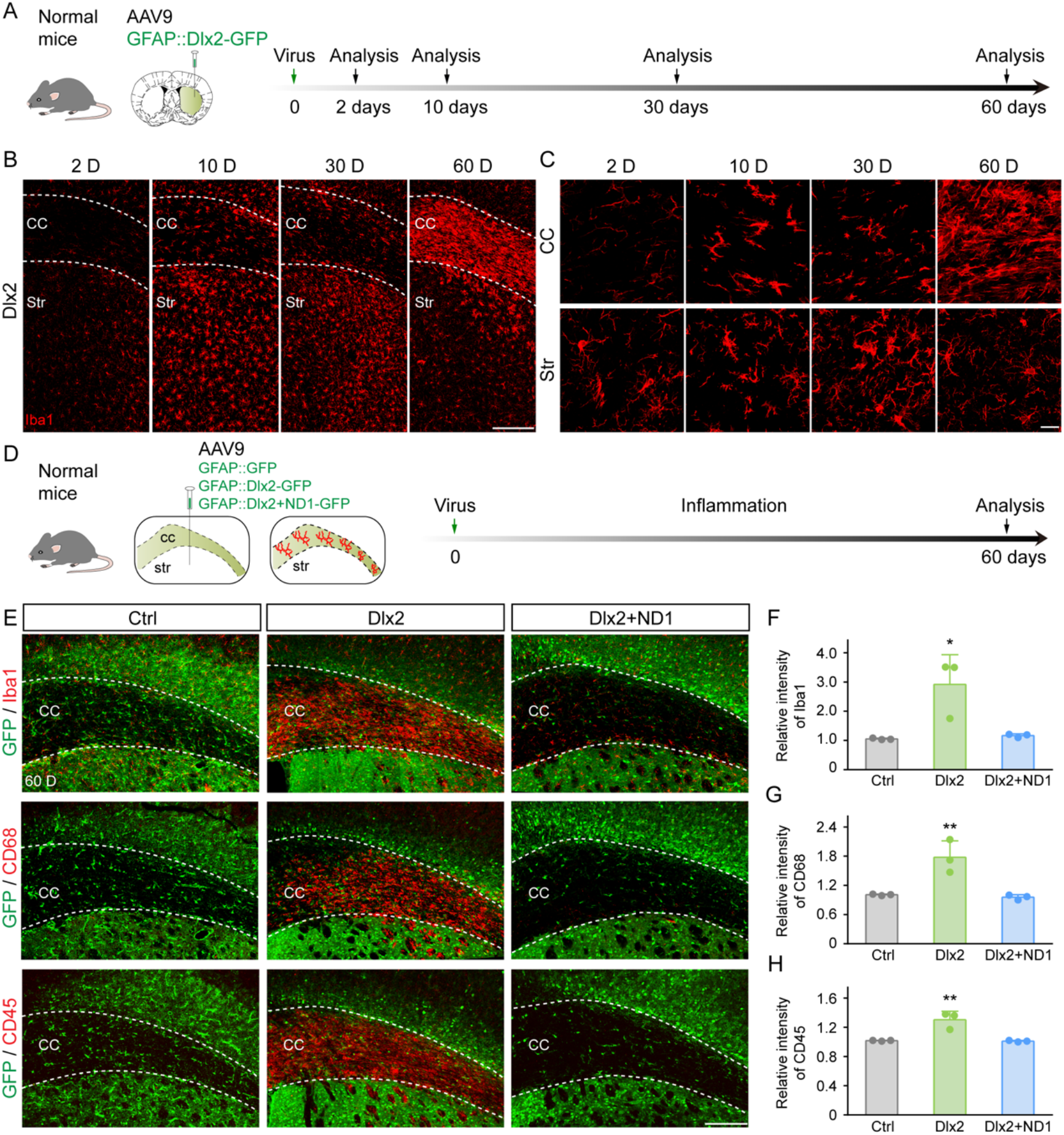
Unexpected Dlx2-induced inflammation in the white matter can be alleviated by NeuroD1. (A) Experimental design to trace microenvironmental change in the infected area of AAV9 GFAP::Dlx2-GFP. (B) Typical images in a series of time points showing Iba1 immunostaining (red) in the corpus callosum (CC) and striatum (Str) following Dlx2-induced AtN conversion. (C) Enlarged images showing different developmental pattern of the Iba1 signal (red) in the corpus callosum (top panel) versus the striatum (bottom panel) following Dlx2 infection. (D) Experimental design to investigate inflammatory response among the three groups of viral infection (control, Dlx2 and Dlx2+ND1). (E) Typical images showing three different inflammatory markers (red; Iba1, top panel; CD68, middle panel; CD45, bottom panel) in the corpus callosum at 60 D after viral infection. Note a significant increase in all three markers in the Dlx2 group (middle panel), which is suppressed in the Dlx2+ND1 group (right panel). (F-H) Quantified data shown in panel E. (Iba1: Ctrl, 1.05 ± 0.03; Dlx2, 2.92 ± 1.01; ND1+Dlx2, 1.17 ± 0.06. CD68: Ctrl, 1.01 ± 0.02; Dlx2, 1.78 ± 0.34; ND1+Dlx2, 0.96 ± 0.05. CD45: Ctrl, 1.02 ± 0.01; Dlx2, 1.30 ± 0.12; ND1+Dlx2, 1.01 ± 0.01. Data presented as mean ± SD. *\*P* < 0.05, *\*\*P* < 0.01, one-way ANOVA with Sidak’s multiple comparison test. n = 3 animals for each group). Scale bars: 200 μm, except C for 20 μm.

### NeuroD1 inhibits Dlx2-induced inflammation in the white matter

We previously did not notice any inflammation in the corpus callosum when we employed NeuroD1 together with Dlx2 to convert striatal astrocytes into neurons ^21^. Therefore, we tested the combination of NeuroD1 and Dlx2 and compared with the effect of Dlx2 alone in the corpus callosum (Fig. 5D for design). Surprisingly, after injecting NeuroD1 together with Dlx2, we found a significant decrease of inflammation in the corpus callosum (Fig. 5E right column, and Fig. S6). Quantified data found that compared to the control group (Fig. 5F-H, gray bars), Dlx2 group showed a significant increase in Iba1, CD68, and CD45 signals in the corpus callosum at 60 days after Dlx2 infection (Fig. 5F-H, green bar), which were significantly attenuated by the addition of NeuroD1 (Fig. 5F-H, blue bar). These results suggest that NeuroD1 can alleviate Dlx2-induced inflammation in the corpus callosum.

We then investigated why NeuroD1 can inhibit the inflammation caused by Dlx2 in the corpus callosum? We previously reported that NeuroD1 did not convert astrocytes in the corpus callosum to neurons nor induced any inflammation in the corpus callosum ^47^. We therefore investigated whether Dlx2 might convert astrocytes in the corpus callosum into neurons. Interestingly, while control AAV GFAP::GFP-infected cells in the corpus callosum showed clear GFAP signal (Fig. 6A, top row, red), many Dlx2-infected cells lost GFAP signal and changed their morphology into round shape but showed no NeuN signal at 60 days after infection (Fig. 6A, middle row). Surprisingly, after adding NeuroD1 together with Dlx2, the viral infected cells showed astrocytic morphology with clear GFAP^+^ signal again (Fig. 6A, bottom row), similar to that found in the control group. Quantitative analysis revealed that compared to the control group (Fig. 6B-C, gray bars), the Dlx2 group showed a significant decrease in the number of GFAP^+^ astrocytes accompanied with a significant increase in the intermediate state cells (GFAP^-^NeuN^-^) in the corpus callosum (Fig. 6B-C, green bars; 60 days after viral infection). Importantly, addition of NeuroD1 reversed such phenomenon: the number of GFAP^+^ cells and the GFAP^-^NeuN^-^ cells both were reversed back to the control level (Fig. 6B-C, blue bars). Similarly, using a different astrocyte marker Sox9, we also found the same reversing effect by NeuroD1, that the loss of Sox9^+^ astrocytes in the Dlx2 group was reversed by the addition of NeuroD1 (Fig. 6D-F). Together, our findings suggest that Dlx2 alone may trigger partial conversion of the astrocytes in the corpus callosum to an intermediate state which is associated with neuroinflammation. Addition of NeuroD1 can reverse such partial conversion and inhibit neuroinflammation caused by Dlx2 alone in the corpus callosum.

**Figure 6.**
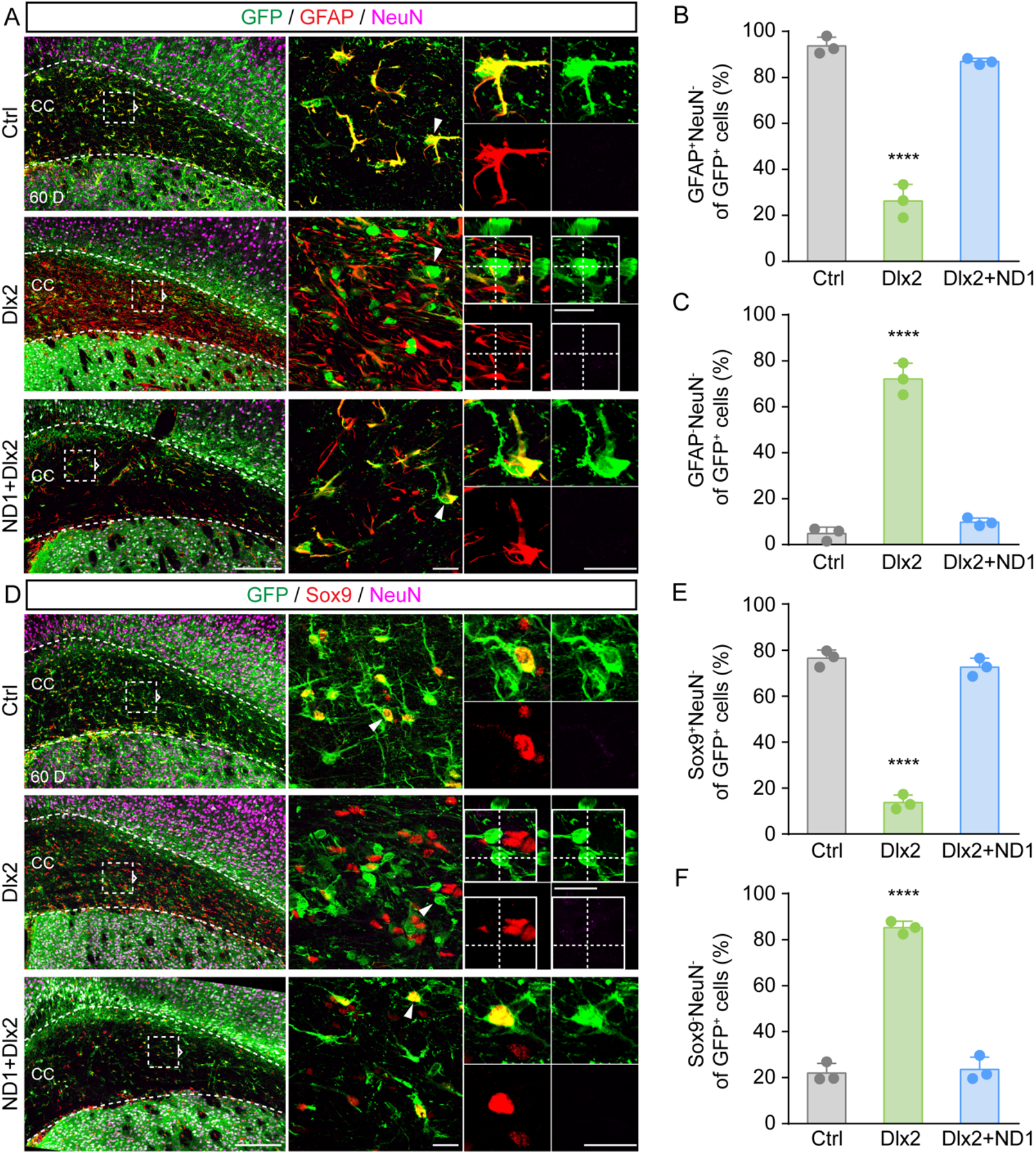
Partial conversion of the astrocytes in the corpus callosum following Dlx2 infection. (A) Typical images showing normal astrocytic morphology with GFAP signal (red) among control GFP-infected (top row) or NeuroD1+Dlx2-infected cells (bottom row) in the corpus callosum. However, the Dlx2-infected cells (middle row) in the corpus callosum lose GFAP signal and become round shape. (B and C) Quantified data of panel A. The percentage of GFAP^+^NeuN^-^ astrocytes among viral infected cells in the corpus callosum is shown in B: Ctrl, 93.7 ± 3.8%; Dlx2, 26.3 ± 7.2%; ND1+Dlx2, 86.9 ± 1.3%. The percentage of GFAP^-^NeuN^-^ intermediate state cells in the corpus callosum is shown in C: Ctrl, 4.8 ± 2.8%; Dlx2, 72.1 ± 6.9%; ND1+Dlx2, 9.8 ± 1.7%. \*\*\*\**P* < 0.0001, one-way ANOVA with Sidak’s multiple comparison test. n = 3 animals for each group. (D) Typical images showing the loss of Sox9 signal among Dlx2-infected cells (middle row) in the corpus callosum, which is also reversed by NeuroD1+Dlx2 (bottom row). (E and F) Quantified data of panel D. The percentage of Sox9^+^NeuN^-^ cells among viral infected cells in the corpus callosum at 60 D is shown in E: Ctrl, 76.5 ± 3.6%; Dlx2, 13.7 ± 3.3%; ND1+Dlx2, 72.6 ± 3.9%). The percentage of Sox9^-^NeuN^-^ cells among viral infected cells in the corpus callosum at 60 D is shown in F: Ctrl, 22.0 ± 4.2%; Dlx2, 85.3 ± 2.7%; ND1+Dlx2, 23.6 ± 5.3%. *\*\*\*\*P* < 0.0001, one-way ANOVA with Sidak’s multiple comparison test. n = 3 animals for each group. Scale bars: 20 μm, except A and D left panel for 200 μm.

## Discussion

This study demonstrates that astrocytes in the adult mouse striatum can be converted into neurons by a single transcriptional factor Dlx2 using different viral systems including retroviruses or AAV. The Dlx2-induced AtN conversion is both dose-dependent and time-dependent, with further confirmation in astrocytic lineage-tracing mice (Aldh1l1::CreER^T2^). During AtN conversion process, there is an intermediate state in-between astrocytes and neurons, where Dlx2-infected astrocytes may lose astrocytic markers but not acquire neuronal markers. Coinciding with astrocyte conversion, there is a transient decrease of astrocytes followed by a quick rebound by an increased proliferation of the remaining astrocytes. An unexpected yet important finding is the activation of microglia in the white matter, including both the corpus callosum and axonal bundles in the striatum, induced by Dlx2 overexpression. One potential cause for such activation of microglia and inflammation is a partial conversion of the white matter astrocytes following Dlx2 infection. Strikingly, addition of NeuroD1 together with Dlx2 inhibits Dlx2-mediated partial conversion in the white matter and consequently alleviates neuroinflammation. These results suggest that Dlx2-induced AtN conversion should be dealt with great caution and adding NeuroD1 together may be one solution to prohibit the inflammation.

### Dlx2-induced striatal astrocyte-to-neuron conversion

We have previously reported that NeuroD1 combined with Dlx2 can efficiently convert striatal astrocytes into medium spiny neurons ^21^. When examining the role of the two transcription factors separately on astrocyte conversion, we discover that Dlx2 alone is sufficient to complete the conversion. Dlx2 belongs to the DLX family of homeobox transcriptional factors and is a key molecular regulator of GABAergic neuron development in the ventral forebrain ^48,49^. Dlx2 alone has been previously demonstrated *in vitro* to convert cultured primary mouse astrocytes into GABAergic neurons ^50^. Dlx2 combined with MicroRNA converts cultured human fibroblasts into medium spiny neurons ^51^. Our data provide further evidence in support of Dlx2 alone to convert striatal astrocytes to neurons *in vivo*. Interestingly, while astrocyte conversion can be achieved through both retroviral and AAV systems, the maturation and survival of the converted neurons are quite different. Most converted neurons induced by retrovirus-Dlx2 are immature neurons with limited numbers, while those converted by AAV-Dlx2 are more mature neurons and can survive much longer. One possibility might be that retrovirus targets proliferative astrocytes that share more stem cell features, whereas AAV can infect both dividing and nondividing astrocytes ^52,53^. Different gene expression and epigenetic profiles of the initial parental cells ^54^ might lead to different progress of AtN conversion. Another possibility might be the differential cellular responses triggered by RNA virus (retrovirus) versus DNA virus (AAV). During the preparation of this manuscript, another paper using lentiviral system also reported the capability of Dlx2 to convert striatal astrocytes into immature neurons ^55^, similar to the immature neurons converted by retrovirus-Dlx2 in our study. Compared to retrovirus and lentivirus, AAV has been approved by FDA as a therapeutic vector and therefore has more translational potential ^56^. Nevertheless, these studies together suggest that AtN conversion triggered by overexpression of neural transcription factors is not restricted to any particular viral system, but rather determined by the expression level of the transcription factors. This is well illustrated by our dose-dependent studies where low dose Dlx2 results in lower conversion rate whereas high dose Dlx2 results in higher conversion rate. On the other hand, the viral dose administered into the brain cannot be too high because any virus injected into the brain will cause anti-viral immune responses, which may result in neural damage ^43^ or unwanted neuronal leakage ^27^. This has also been nicely illustrated in our dose-finding studies using control AAV9 GFAP::GFP, where high titre at 1E13 GC/ml results in severe activation of microglia. In our most recent study, we show that injection of high titre 1E13 GC/ml AAV9 GFAP::GFP (1 μl) results in 50-60% neuronal leakage ^37^, highlighting the necessity to use low titre AAV for conversion studies. Based on this study and our previous studies, we recommend that AAV titre should be kept in a safe range of ~E11 to E12 GC/ml when conducting AtN conversion studies. Previous studies using high titre of AAV (>E13 GC/ml) such as that by Wang et al ^27^ will inevitably have high neuronal leakage and should be repeated with lower titre AAV to validate their results.

### Dlx2 changes the astrocyte identity in the corpus callosum and induces inflammation

We have previously reported that NeuroD1 efficiently converts astrocytes in the grey matter such as the cortex into neurons, but not the astrocytes in the white matter such as the corpus callosum ^47^. Similarly, combination of neurogenin-2 and Nurr1 together has also been reported to convert cortical astrocytes into neurons but not the astrocytes in the corpus callosum ^13^. The fibrotic astrocytes in the white matter have different morphology and gene expression profile from the protoplasmic astrocytes in the grey matter ^57^, which may contribute to their differential response to the ectopically expressed neural transcription factors. In addition, white matter is composed of myelinated axons with very few neurons, which may not support the survival of newly converted neurons if any. Different from the effect of NeuroD1 or Neurogenin2+Nurr1 in the white matter, this study discovers a surprising capability of Dlx2 in changing the identity of astrocytes in the corpus callosum. The Dlx2-infected astrocytes in the corpus callosum lose astrocytic markers such as GFAP and Sox9 but do not acquire neuronal markers such as NeuN. Such unique GFAP^-^Sox9^-^NeuN^-^ property suggests that overexpression of Dlx2 in the white matter astrocytes might have turned them into an intermediate state that are neither astrocytes nor neurons and consequently trigger the activation of microglia to engage them. In fact, consistent with our finding, careful analysis of the recent work using lentivirus-Dlx2 to convert striatal astrocytes also revealed activation of microglia in the Dlx2-injected side (Zhang et al., 2022; their Suppl Fig. S3A-B, middle row), although the authors were not aware of such drastic difference in their Iba1 staining. It is also worth to point out that in the lentivirus-Dlx2 conversion paper^55^, there appears to be a severe neuronal loss as shown by only a few NeuN^+^cells in their Fig. 3D-G. While the authors themselves may have overlooked this important neuronal loss, it might be linked to their overlooked inflammation as well. Our discovery of the neuroinflammation associated with Dlx2-conversion raises serious concerns regarding its therapeutic potential if Dlx2 is applied alone. In general, we advocate that great caution should be taken to safeguard all the applications involving AtN conversion technology for therapeutic purposes.

### Unexpected anti-inflammation role of NeuroD1 after combined with Dlx2

While it is unexpected that Dlx2-induced AtN conversion would result in neuroinflammation, it is equally unexpected that adding NeuroD1 would suppress the inflammation induced by Dlx2. The Dlx2-induced inflammation appeared to be associated with a partial conversion induced by Dlx2 in the white matter astrocytes. Interestingly, adding NeuroD1 together with Dlx2 prevents such partial conversion in the white matter, similar to our previous finding that NeuroD1 alone cannot convert white matter astrocytes into neurons ^47^. Why NeuroD1 cannot convert white matter astrocytes yet Dlx2 can partially convert white matter astrocytes into an intermediate state is an outstanding question that awaits further investigation. On the other hand, this is not the first time we observe an anti-inflammation role of NeuroD1 during AtN conversion. We have previously reported that overexpression of NeuroD1 in reactive astrocytes after stab injury results in a significant reduction of A1 astrocytes, accompanied with a decrease of neuroinflammation ^31^. In addition, in an ischemic cortical stroke model, we have also reported that after NeuroD1-mediated AtN conversion, the microglia-associated inflammation has been significantly attenuated as well ^20^. Together, these studies support an important role of NeuroD1 in anti-inflammation that may be beyond its role in promoting AtN conversion.

### Perspective on AtN conversion and its potential application

This study, for the first time, raises a serious concern that AtN conversion might result in an unexpected side effect, such as neuroinflammation in the white matter. It highlights the fact that our understanding of *in vivo* AtN conversion is still in early stage with many more to be explored. On the other hand, such potential side effect should not be viewed as a roadblock that will deter our advancement toward future clinical applications. The fact that addition of NeuroD1 can overcome the Dlx2-induced inflammation is a good example that side effects can be mitigated if we find a solution. The sharp contrast in the effect of NeuroD1 versus Dlx2 on the astrocytes of the white matter also underscore the importance of testing the right factors in the right place. Generating ectopic neurons in the wrong place, such as generating neurons in the white matter or generating dopaminergic neurons in the striatum instead of the substantial nigra, might result in unwanted consequences such as forming wrong neural circuits. An ideal strategy for AtN conversion is using local glial cells to generate local neurons without obvious side effects in the grey matter and the white matter.

## Materials and methods

### Animals

Both adult male and female wild-type C57BL/6J mice (Guangdong Medical Laboratory Animal Center) and Aldh1l1::CreER^T2^ transgenic mice (The Jackson Laboratory, RRID:IMSR_JAX:031008) at 2-5 months of age were recruited in the experiments. Mice were housed in a 12 h light/dark cycle with food and water *ad libitum*. All experimental protocols were approved by the Jinan University Institutional Animal Care and Use Committee (approval No. IACUC-20180321-03).

### Stab injury and viral injection

Adult wild-type C57BL/6J mice were used for stab injury as described ^31^ with slight modification. Briefly, animals were anesthetized by intraperitoneal injection of 20 mg/kg Avertin (a mixture of 12.5 mg/mL of 2,2,2-Tribromoethanol and 25 μl/mL 2-Methyl-2-butanol) and placed in a stereotaxic apparatus with the skull exposed. Holes were drilled in bilateral skull above the striatum (Coordination: anterior/posterior (AP): +1.0 mm; medial/lateral (ML): +/-2.0 mm relative to Bregma). A blunt needle (~1.0 mm) placed into each site to the depth of −3.5 mm dorsal-ventral (DV) into striatum and stayed still for 3 min. Three days later when striatal glia already became reactive, 2 μl retrovirus (CAG::GFP-IRES-GFP / CAG::Dlx2-IRES-GFP) were injected into the same site. For AAV9 administration, 2 μl virus was injected into right intact striatum Aldh1l1::CreER^T2^ mice or wild-type C57BL/6J mice at the following coordination: AP: 1.0 mm, ML 2.0 mm relative to Bregma, and DV 3.0 mm relative to the dura. The virus injection was performed using a 5 μl micro-syringe with a glass pipette at the speed of 200 nl/min. After injection, the pipette was kept in place for about 10 min.

### Tamoxifen administration

Tamoxifen (Sigma, T5648) was dissolved in corn oil at the concentration of 15 mg/mL. Tamoxifen was administrated through intraperitoneal injection at a daily dose 1 mg/10 g body weight for 3-5 consecutive days after virus injection.

### Virus information

Retroviral vectors were constructed, packaged and concentrated according to the described ^20^. *hGFAP* promoter used in the AAV vectors is the synthetic 681-bp gfaABC1D. The construction of *pAAV GFAP::GFP, GFAP::Dlx2-P2A-GFP, GFAP::NeuroD1-P2A-GFP*, the following AAV9 virus package, purification with iodixanol and subsequent concentration, as well as virus titration were produced by PackGene^®^ Biotech, LLC (Guangzhou, China). All virus used in this study was diluted and stocked in 0.001% Pluronic F-68 solution (Poloxamer 188 Solution, PFL01-100ML, Caisson Laboratories, Smithfield, UT, USA).

### Tissue preparation and immunofluorescence

Animals were deeply anesthetized and then performed intracardiac perfusion with 0.9% saline and subsequent ice-cold 4% PFA. The brain tissues were collected and post-fixed overnight in 4% PFA at 4°C. Then they were placed in a series of 10%, 20%, 30%, and 30% sucrose until the tissue sank. After the embedment in Optimal Cutting Temperature (Tissue-Tek^®^ O.C.T.), brain tissues were frozen at −80°C. The cerebral regions of interest were serially sectioned using the cryostat (Thermo Scientific) at 30 μm thickness. For immunostaining, the brain slices were firstly washed 3 times in phosphate buffered saline (PBS, pH: 7.35, OSM: 300) for 5 min each, and then blocked in 0.3% Triton X-100 + 5% normal donkey serum (NDS) + 5% normal bovine serum albumin (BSA) for 1 hour. The primary antibodies prepared in this blocking solution were added and incubated at 4°C for 1-2 night. After 30 min recovery at room temperature, the brain slices were washed 3 times in PBST (PBST+0.2% Tween 20) for 15 min each. Afterwards, the samples were incubated with corresponding secondary antibody conjugated to Alexa Flour 448, Alexa Flour 555 or Alexa Flour 647 (1:1000 except Alexa Flour 647 for 1:500; Life technologies) and 4’,6-diamidino-2-phenylindole (DAPI for nuclei counterstaining, 1:2000) for 2 hours at room temperature and subsequent wash with extensive PBS. Later, brain slices were carefully mounted with anti-fade mounting medium (VECTOR). All representative images were captured with a Zeiss confocal microscope (LSM 880) at 200x, 400x and 630x magnifications or a Carl Zeiss Microscopy (Imager Z2) at 100x magnification.

### TUNEL staining

Brain slices were washed 2 times in PBS for 5-10 min each, and then incubated in PBS with 0.5% Triton X-100 for 5 min. The terminal deoxynucleotidyl transferase (TdT) buffer (One Step TUNEL Apoptosis Assay Kit, Beyotime, China) containing TdT (1:10) and Cy3-labeled dUTP was added and incubated for 1 hr at 37°C. Subsequently, the brain slices were immunostained with other antibodies.

### Quantification and statistical analysis

All images with the same antigen were captured and adjusted with the same parameters. For quantification, 24 fields (0.0108 mm^3^) at the 400x magnification (Fig 1D and 1G; Fig 2C, 2F and 2H; Fig 3E-F; Fig 4C; Fig 5B-C and 5E-F) and 3 fields (1.1213 mm^2^) at the 100x magnification (Fig 4E-F) in the striatum were randomly chosen from 3 brain slices. The relative intensity of Iba1, CD68 and CD45 from virus side of the corpus callosum were acquired using those from contralateral side as an index (Fig 5F-H). The density of Sox9^+^ cells was measured with Image J software. Other images were analyzed with Zeiss ZEN 2.3 software (blue edition).

All Statistical analyses were performed double-blindly. Quantification data were shown as mean or mean ± standard deviation (SD). Statistical analysis was performed by two-tailed Mann-Whitney test (unpaired) for two-group comparison, one-way ANOVA with Sidak’s multiple comparison test for multiple group comparisons, and two-way ANOVA analysis followed by Sidak’s multiple comparison test for multivariate and multiple group comparisons using GraphPad Prism software v.8.0. A *P* < 0.05 was considered significant.

## Acknowledgments

This study was supported by the National Natural Science Foundation of China, Nos. U1801681 (to G.C.), 31701291 (to W.L.) and 32100793 (to Z.-Q.X.), Foundation of Guangdong Province of China, No. 2021A1515011237 (to W.L.), and the Guangdong Province Science and Technology Planning Project of China, No. 2018B030332001 (to G.C.). It was also supported by Yi-Liang Liu Endowment Fund from Jinan University Education Development Foundation. We would like to thank all the Chen lab members from Guangdong-Hong Kong-Macau Institute of CNS Regeneration (GHMICR) at Jinan University for rigorous discussion and valuable suggestion.

## Author contributions

G.C., W.L., M.-H.L. and Y.-G.X. designed most of the experiments, analyzed the data, and wrote the manuscript. M.-H.L. and Y.-G.X. carried out most of the experiments in mice, quantified the data, and made the figures. W.L. provided the retrovirus for this study. X.-N.B., J.-H.L., Z.-Q.X., T.W., and L.X. all contributed to these studies including immunostaining and quantification.

## Supplementary Material

**Supplementary Figure 1.**
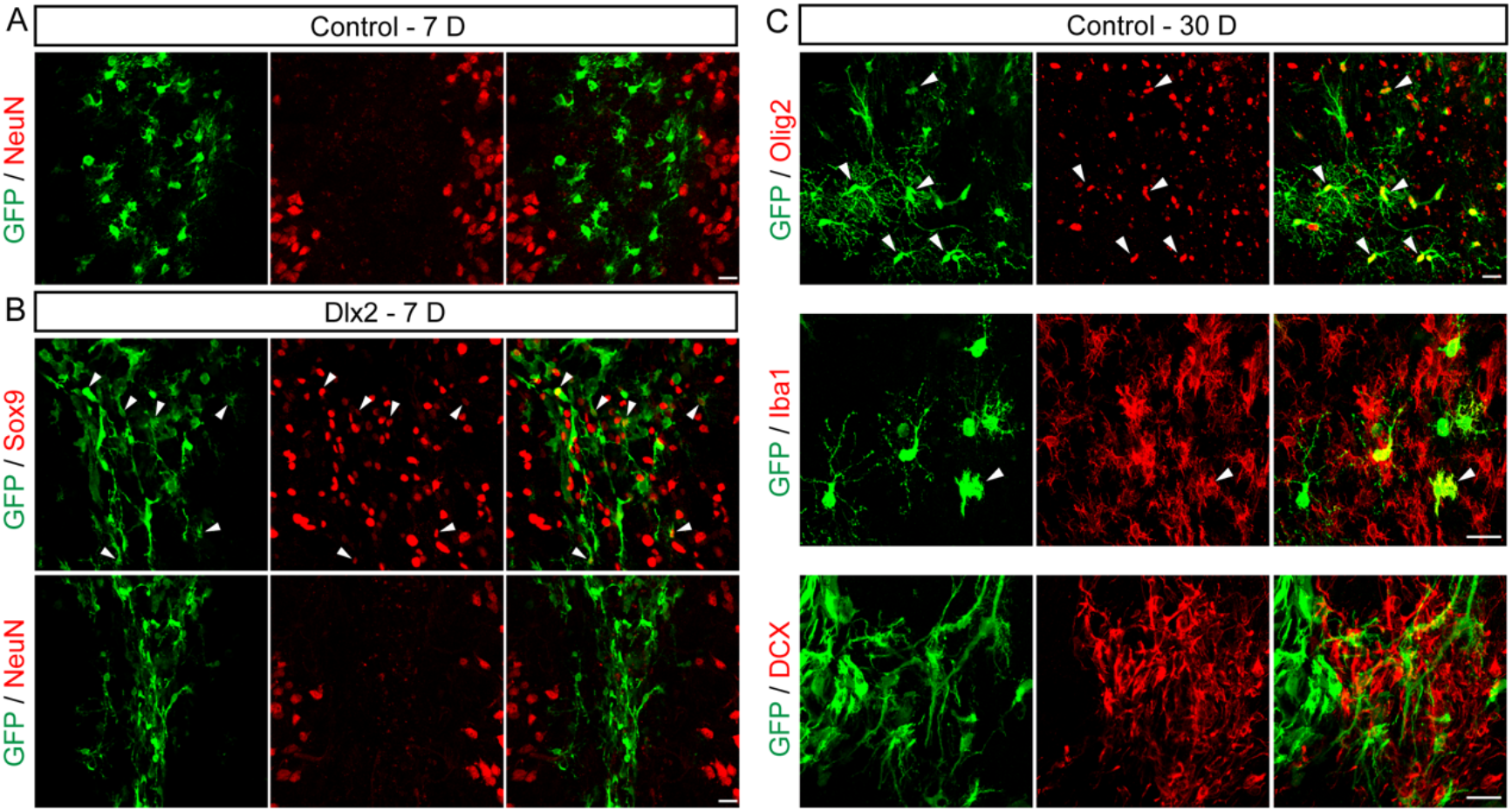
Retrovirus specifically expresses transgenes in glial cells after stab injury. (A) None of the GFP^+^ cells (green) show NeuN signal (red) in the control group at 7 days post viral infection (7 D). (B) Some of the Dlx2-GFP^+^ cells (green) expressed Sox9 (red, top row), but NeuN signal (red, bottom row) is rarely detected in the Dlx2 group at this early stage of 7 D. (C) Representative images at high magnification of GFP^+^ cells coimmunostaining with different markers in the control group at 30 D (Olig2, top row; Iba1, middle row; DCX, bottom row). Scale bars: 20 μm.

**Supplementary Figure 2.**
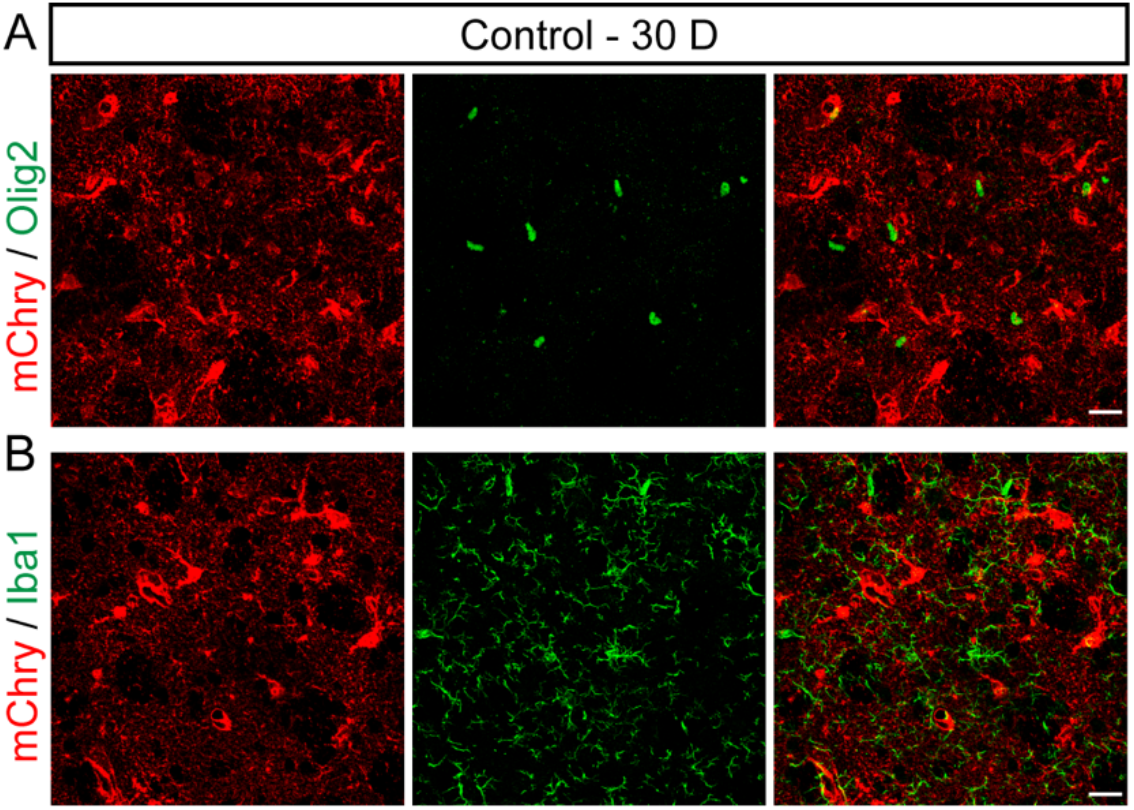
AAV9 FLEX system doesn’t target striatal oligodendrocyte and microglia in Aldh1L1::CreER^T2^ mice. (A and B) Representative images show that Olig2 signal (green, A) and Iba1 signal (green, B) are rarely detected in mCherry^+^ cells (red) at 30 D. Scale bars: 20 μm.

**Supplementary Figure 3.**
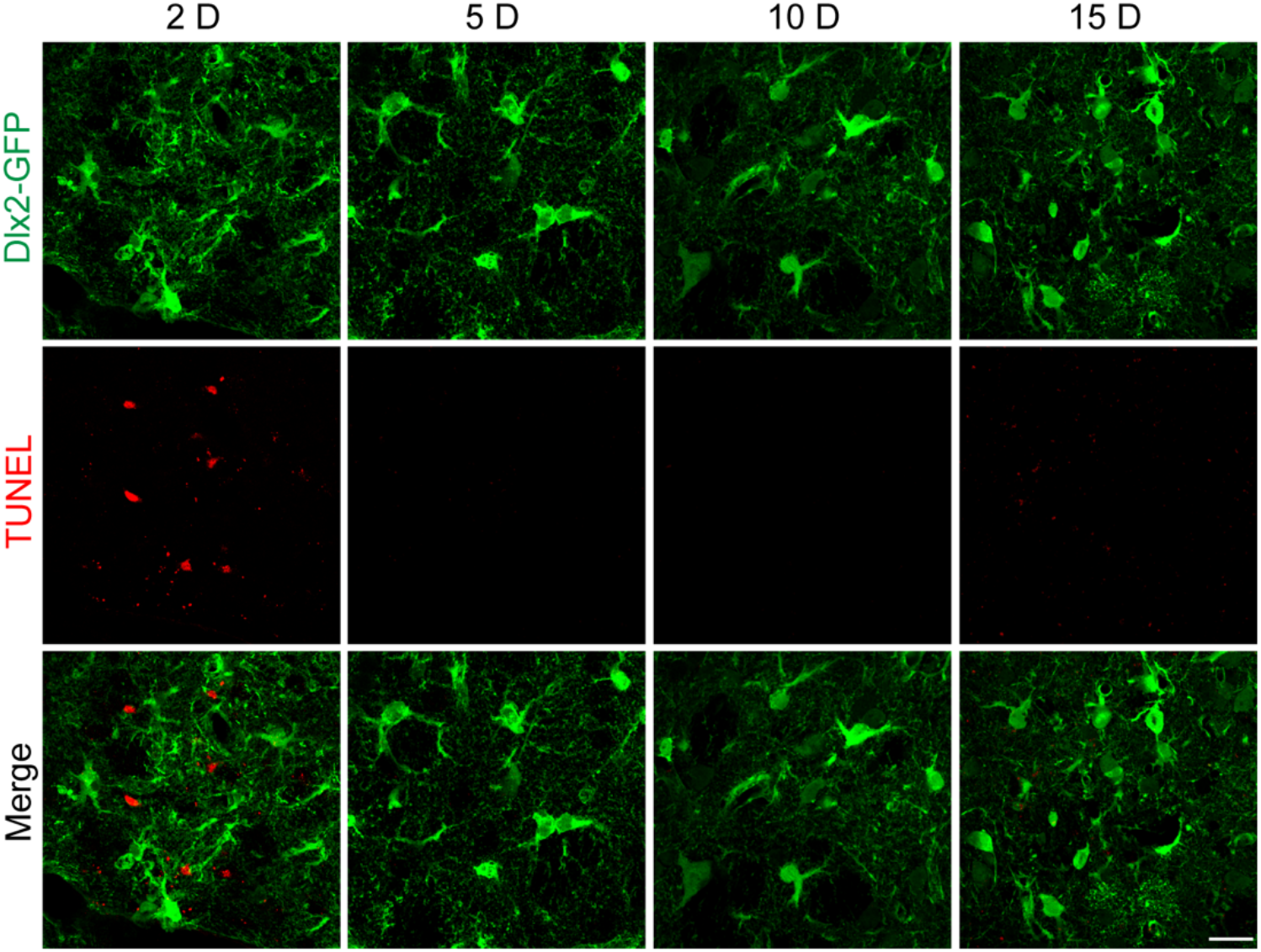
Apoptosis is rarely detected in Dlx2-converted cells. During the conversion process (5-15 dpi), TUNEL signal (red) was rarely detected in the Dlx2-GFP^+^ cells, while a few cells adjacent to the needle track were positive for TUNEL staining suggesting stab injury-caused cell apoptosis at 2 days post viral injection.

**Supplementary Figure 4.**
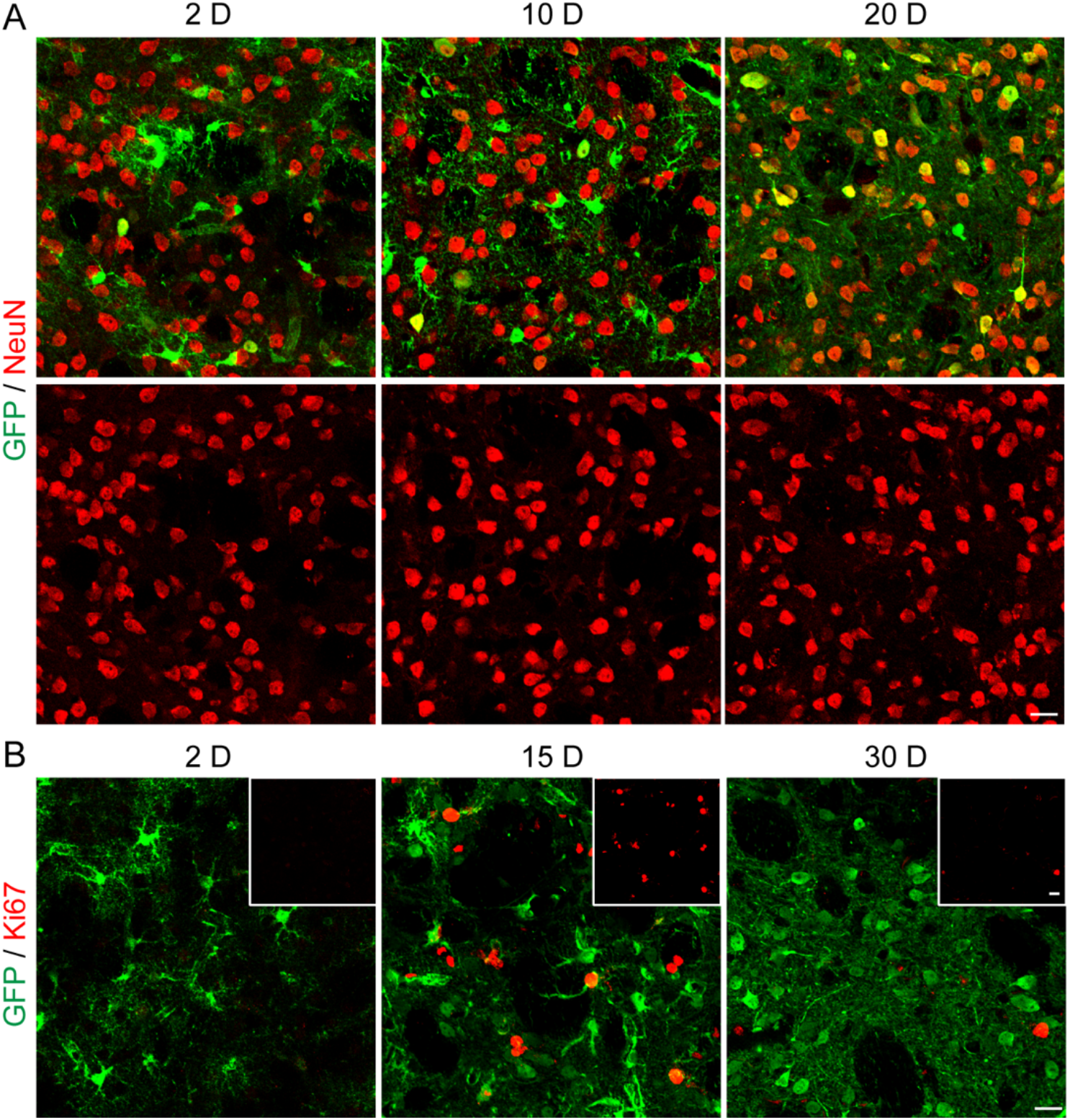
Increased astrocytic proliferation following astrocyte-to-neuron conversion in the striatum. (A) Representative images showing no adversary effect of Dlx2-induced astrocyte-to-neuron conversion on NeuN signal despite a transient decrease of Sox9 signal among Dlx2-infected cells around 10 D. (B) Representative images showing an increase of Ki67^+^ cells (red) within the Dlx2-infected areas (GFP, green) at ~15D in the mouse striatum. Scale bars: 20 μm.

**Supplementary Figure 5.**
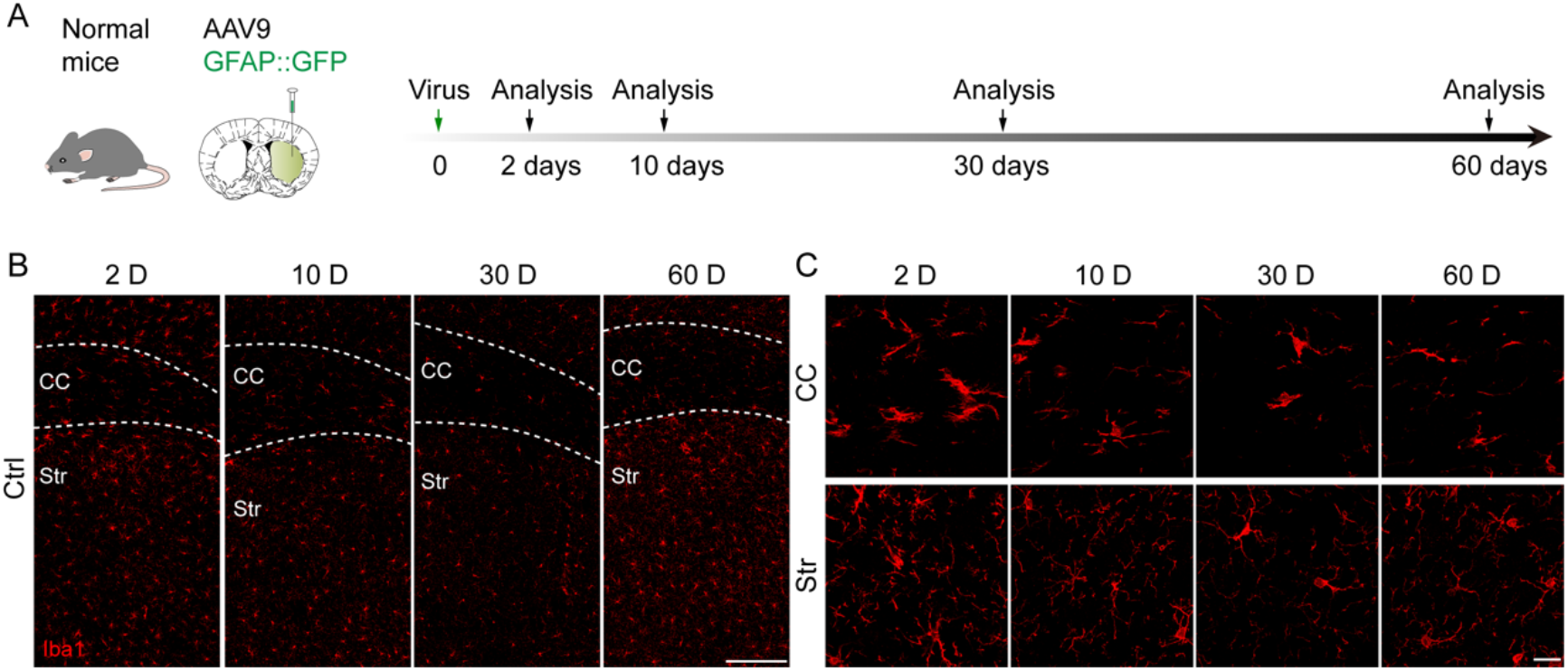
AAV9 GFAP::GFP doesn’t cause inflammation in the white matter and striatum at proper titer. (A) Experimental design to trace microenvironmental change in the infected areas by control AAV9 GFAP::GFP. (B and C) Typical images in a series of time points showing Iba1 immunostaining (red) in the corpus callosum (CC, top panel) and striatum (Str, bottom panel) of control virus infection. Scale bars: 200 μm for B, 20 μm for C.

**Supplementary Figure 6.**
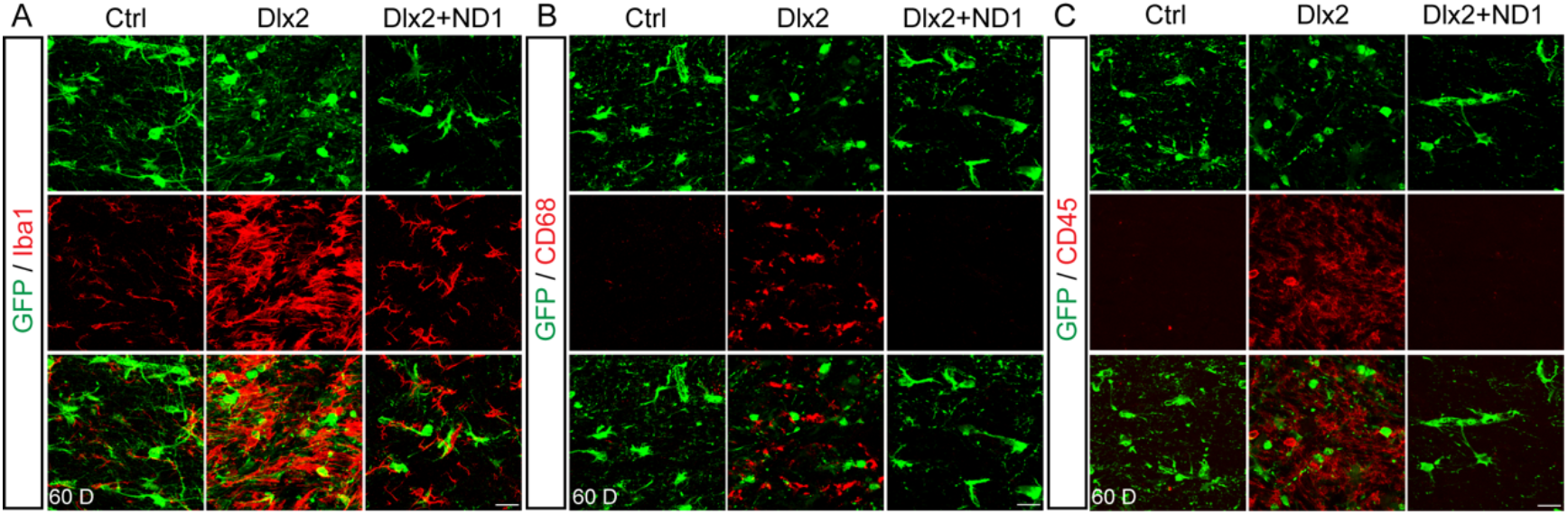
Dlx2 causes inflammation in the corpus callosum. (A-C) Representative images at high magnification showing inflammatory markers (Iba1, A; CD68, B; CD45, C) in three groups (control, left panel; Dlx2, middle panel; Dlx2+ND1, right panel). There is a significant increase in all three inflammatory markers in the corpus callosum of the Dlx2 group at 60 D. Scale bars: 20 μm

**Supplementary Table1.** Virus

| **Virus type** | **Use titers** |
| --- | --- |
| Retrovirus ***CAG::GFP-IRES-GFP*** | 1E8-5E8 TU/mL |
| Retrovirus ***CAG::Dlx2-IRES-GFP*** | 1E8-5E8 TU/mL |
| AAV9 ***Flex-CAG::mCherry-P2A-mCherry*** | E12 GC/mL |
| AAV9 ***Flex-CAG::Dlx2-P2A-mCherry*** | E12 GC/mL |
| AAV9 ***GFAP::GFP*** | 2E10, 2E11, 2E12, 1E13<br>GC/mL |
| AAV9 ***GFAP::Dlx2-P2A-GFP*** | 2E10, 2E11, 2E12 GC/mL |
| AAV9 ***GFAP::NeuroD1-P2A-GFP*** | 2E10, 2E11, 2E12 GC/mL |

**Supplementary Table2.** Antibodies

| **Reagent** | **Source** | **Identifier** | **Dilution** |
| --- | --- | --- | --- |
| Rat anti-Ctip2 | Abcam | Ab18465 | 1:500 |
| Rat anti-CD45 | BD Pharmingen | 550539 | 1:500 |
| Rat anti-CD68 | Abcam | Ab125212 | 1:500 |
| Rabbit anti-DARPP32 | CST | 2306S | 1:1000 |
| Rabbit anti-DCX | Abcam | Ab18723 | 1:1000 |
| Guinea pig anti-DCX | Millipore | Ab2253 | 1:2000 |
| Rabbit anti-Dlx2 | Abcam | Ab135620 | 1:500 |
| Rabbit anti-GFAP | Sigma | G9269 | 1:1000 |
| Rat anti-GFAP | Invitrogen | 13-0300 | 1:1000 |
| Chicken anti-GFP | Abcam | Ab13970 | 1:2000 |
| Rabbit anti-Iba1 | Wako | 019-19741 | 1:500 |
| Rabbit anti-Ki67 | Abcam | Ab15580 | 1:500 |
| Chicken anti-mCherry | Abcam | Ab205402 | 1:1000 |
| Mouse anti-NeuN | Novus | NBP1-92693 | 1:300 |
| Rabbit anti-NeuN | Abcam | Ab177487 | 1:1000 |
| Guinea pig anti-NeuN | Millipore | ABN90 | 1:1000 |
| Rabbit anti-NeuroD1 | Abcam | Ab205300 | 1:500 |
| Rabbit anti-Olig2 | Millipore | AB9610 | 1:500 |
| Rabbit anti-Sox9 | Millipore | AB5535 | 1:500 |
| Rabbit anti-S100β | Abcam | Ab52642 | 1:1000 |
| Chicken anti-MAP2 | Abcam | Ab5392 | 1:2000 |

